# Population dynamics in pre-Inca human groups from the Osmore Valley, the Azapa Valley and the coast of the south central Andes

**DOI:** 10.1101/2020.02.06.936823

**Authors:** A. Coppa, F. Candilio, C. Arganini, E. de la Vega, E.G. Moreno Terrazas, M. Lucci, A. Cucina

## Abstract

The present study applies a dental morphological perspective to the understanding of the complex population history of pre-contact South-Central Andes, the detection of the underlying dynamics, and the assessment of the biological ties among groups. It takes into account 1665 individuals from 16 sites that date from the Archaic to the Late Intermediate located along the coast, on the *altiplano*, and in the coastal valleys of both Chile and Peru. The results obtained highlight the need for wider perspectives capable of taking into account both the different micro-regional realities and the region in its ensemble. The population dynamics and mobility patterns detected indicate the widely accepted interpretations and distinctions based on cultural affiliations might be insufficient to comprehend the complex population history of the region, especially because the results obtained in the present study indicate the presence of a general and widespread common morphological background for the inhabitants of some of these cultures (i.e., Moche and Wari) and that the interactions they had throughout time must have been far from inconsequential.

## Introduction

The South Central Andes extend over a vast territory that encompasses southern Peru and northern Chile; an area that is of particular bio-archaeological significance in light of the different cultures that have settled there throughout time and the complex population dynamics that have characterized settlement patterns and migrations. Archaeological evidence has enabled the detection of an important change in subsistence between 8,000 and 1,000 BC, in which the egalitarian maritime populations with fishing economies attested as of 12,000 BC [1] gradually adopted production systems of subsistence [2]. The development of more centralized forms of power led to the rise of important empires – such as the Moche, the Wari and the Tiwanaku – that extended their authority through a variety of different strategies: colonization by agropastoral migration in the case of Tiwanaku and centralized expansion in that of Wari [3]. Though consensus has yet to be reached regarding these migrations and their influence on the populations that inhabited the region, there has been intense academic debate that has also resulted in the formulation of different hypotheses regarding the entity, the extent, and the impact of the Tiwanaku colonies in various valleys in the region. One may, however, state that – in general – Tiwanaku was likely founded in the southern Titicaca Basin, immediately south of Lake Titicaca, during the first or second century AD and that it thrived becoming the capital of an archaic state from around AD 500-1000 [4]. Tiwanaku reached its apogee during the Middle Horizon period, around AD 700 [5]), and began its decline by AD 900 [6]. Archaeological evidence suggests it expanded its sphere of influence by establishing permanent colonies in many surrounding territories [3,7] among which the Osmore – or Moquegua – Valley in southern Peru [8] and the Azapa Valley in northern Chile [9-14].

The growing number of human collections analyzed from pre-contact period in the region has led to extensive investigations of the cultural, genetic and ethnic changes that took place in these valleys in pre-Inca times [5,15-24]. According to Varela and Cocilovo [13], the Azapa Valley was originally inhabited by Archaic fishermen and experienced the arrival of settlers likely from the circum-Titicaca area during the Early Intermediate (Formative) period. Though the authors state the migrations appeared to have had greater impact on the populations that lived in the valleys compared to those that lived on the coast, they believe the substantial gene flow between these two regions created a clear similarity between these two groups. They hypothesize that, in part due to the strong influence exerted on the valley by the Tiwanaku, as of AD 300, these two populations started to change and diversify. The model depicted in their study indicates that there must have been an increase of regional gene flow to the valley throughout the Tiwanaku period (AD 300-800) and that this, in combination with the gradual reproductive isolation of the coastal populations, brought to the progressive differentiation of these previously very similar groups. It is only during the Late Intermediate – after the decline of the Tiwanaku Empire between AD 800-1,000 and possibly in consequence to it – that they detect a substantial decrease of gene flow to the valley. On the contrary, Sutter and Mertz [24] believe there is no direct evidence indicating the Tiwanaku replaced the Azapa groups during the Middle Horizon. Their study indicates there is continuity through time in the valley and suggests that the similarities observed with the Tiwanaku are, most likely, the effect of interaction and cultural exchange rather than the product of intense immigration into the valley from the northeastern regions south of the Titicaca Basin. In a later study, Sutter [22] detected biological continuity among coastal groups as well as among inland groups in the Azapa Valley that was not accompanied by significant, if any, continuity between these two regions; thus further supporting his theories that refute significant influence of allochtonous populations to the local gene pool.

The most widely accepted hypotheses regarding the population history of the Osmore Valley indicate that ethnically diverse colonists from the Tiwanaku empire spread and settled into different areas of the Upper Valley, where they established permanent outposts such as Chen Chen and Omo [3,25-26]. Archaeological evidence [3] suggests they did so through a variety of settlement patterns and that, in the Osmore Valley, the Tiwanaku expansion proceeded at a massive demographic scale and through direct colonization. According to this hypothesis, the colonists settled into four different enclaves, avoided transculturation with the valley’s indigenous inhabitants [17,25] and continued to receive a small influx of individuals from different parts of the Tiwanaku polity through time [27]. Regardless of the extremely strong influence it had on the valley, Tiwanaku does not appear to have ever obtained complete territorial control of it but, instead, to have coexisted, presumably peacefully [26], with both the indigenous inhabitants and, as of AD 600, with Wari outposts such as Cerro Baúl [28-30]. These studies indicate that regardless of the close proximity (at times of no more than 10-20 km) and the long period of time throughout which they must have necessarily interacted, intercultural transmission in the valley was extremely limited and each of these communities maintained its own traditional customs, architectures and technologies [25].

Even though widely accepted, the abovementioned hypotheses regarding the population history of the Osmore Valley are not the only ones that have been proposed. In fact, Mosely et al. [31] believe the Tiwanaku population in the Osmore Valley could have been an autonomous group that, under the Tiwanaku sphere of influence [15], adopted by means of cultural transmission the Tiwanaku customs and way of life for, as stated by Blom, “…the mere possession of these (Tiwanaku) artifacts might not indicate population movement and shifts in identity.” [15: p.155].

Given that the presence of shared material culture may indirectly suggest biological contact between different entities or groups but it may not demonstrate it or provide direct evidence of population admixture, the studies have, in the past years, shifted towards multidisciplinary approaches that include bio-archaeological analyses based on evolutionary theory. These have provided great insight into the population dynamics of the region but have, to this day, mainly focused on specific research questions or areas [15-17,24,26,32-36]. Exceptions to this are a number of studies conducted by Sutter [2,5,22] that widen the perspective by taking into account a vaster area that includes both the Osmore and the Azapa Valleys. These studies analyze, through different means, the complex settlement histories that characterize each of the valleys in order to better understand the region as a whole as well as the ties between the different populations that inhabited it. The results indicate that the situations depicted in the different areas are far from homogeneous and that while there are indications that the Late Intermediate inhabitants of the Osmore Valley may derive from immigrant populations, the ones in the Azapa Valley appear to derive from in situ microevolution. These differences certainly indicate much is yet to be understood regarding the population history and the biological ties between the different groups that inhabited both valleys and the *altiplano* [37] and reaffirm the necessity of large-scale multidisciplinary approaches.

Given the centrality of the Tiwanaku to the research question we believe the area under investigation needs to be broadened even further to include the southern portion of the Titicaca basin and address true Tiwanaku samples and not solely their supposed outposts to the valleys. We feel this is especially crucial given the results obtained by us in a previous study [18] that suggest caution in considering the inhabitants of the outposts to the valleys representative of the presumed ancestral population, especially when dealing with complex peopling scenarios as is the case in the South Central Andes. The study, a smaller scale analysis of the pre-Inca inhabitants of the Osmore Valley conducted using dental morphological traits, compared Formative, Middle Horizon and Late Intermediate samples among which the supposed Tiwanaku outpost of Chen Chen. It showed affinities between the Chen Chen and the Wari samples suggesting that either the theories according to which the inhabitants of the Tiwanaku colonies avoided transculturation with the autochthonous groups should be revisited or that perhaps Chen Chen’s attribution as a Tiwanaku colony itself should be questioned.

The present study wishes to address a number of research questions focused on the Tiwanaku, the manner in which they colonized new territories, the ties they established with the neighboring communities and those that connected them to the inhabitants that preceded and followed them in the region. It does so by including in the study data pertaining to a small but central sample formed of 35 Middle Horizon individuals from Tiwanaku as well as from 119 Late Intermediate individuals from the circum-Titicaca region – thus availing itself of direct evidence in place of a proxy such as Chen Chen – and addressing the complex debates regarding continuity versus replacement and in-situ biological change versus migration.

## Materials and methods

The present analysis comprises dental morphological data pertaining to 16 Peruvian, Bolivian and Chilean pre-Inca sites (Fig 1) (see also S1 Table), which range from the Archaic to the Late Intermediate period, with a total of 1629 dentitions (Table 1 and see Table S1). The data pertaining to the Formative site of Moche, the Middle Horizon sites of Chen Chen, Wari and Tiwanaku, and the Late Intermediate sites of Chancay and Titicaca were scored directly by one of the authors of the present study (Arganini, Candilio or Coppa), whereas the data pertaining to the Late intermediate sites of Chiribaya, Yaral and San Geronimo as well as to the Chilean sites from the Azapa Valley were obtained from the literature [21]. Tables S2 to S17 show the individual data of all the 79 morphological traits scored in each of the 16 groups that have been analyzed in this paper. They also include data recovered form the literature [21] for comparative purposes.

**Table 1.** List of sites analyzed, acronym, chronological period and geographic location.

|  | Site | Acronym | Sample size | Period | Chronology | Region |
| --- | --- | --- | --- | --- | --- | --- |
| 1 | ARCHAIC_CHILE | ARL | 84 | Archaic | 5000-1000 BC | Northern Chile |
| 2 | ARCHAIC_PERU | ARP | 16 | Archaic | 5000-1000 BC | Southern Peru |
| 3 | AZAPA_COST_FO | AFC | 71 | Formative | 1000 BC – AD 200 | Northern Chile |
| 4 | AZAPA_VALLEY_FO | AFV | 62 | Formative | 1000 BC – AD 200 | Northern Chile |
| 5 | MOCHE_FO | MFO | 157 | Formative | 1000 BC – AD 200 | Northern Peru |
| 6 | AZAPA_MH | AMH | 191 | Middle Horizon | AD 200-900 | Northern Chile |
| 7 | TIWANAKU_MH | TMH | 35 | Middle Horizon | AD 200-900 | Northern Bolivia |
| 8 | WARI_MH | WMH | 65 | Middle Horizon | AD 200-900 | Northern Peru |
| 9 | AZAPA_COAST_LI | ALC | 43 | Late Intermediate | AD 900-1450 | Northern Chile |
| 10 | AZAPA_VALLEY_LI | ALV | 25 | Late Intermediate | AD 900-1450 | Northern Chile |
| 11 | CHIRIBAYA_LI | CIR | 185 | Late Intermediate | AD 900-1450 | Southern Peru |
| 12 | SAN_GERONIMO_LI | SGE | 58 | Late Intermediate | AD 900-1450 | Southern Peru |
| 13 | YARAL_LI | YAR | 52 | Late Intermediate | AD 900-1450 | Southern Peru |
| 14 | CHEN_CHEN_LI | CHE | 119 | Late Intermediate | AD 900-1450 | Southern Peru |
| 15 | TITICACA_LI | TLI | 195 | Late Intermediate | AD 900-1450 | Southern Peru |
| 16 | CHANCAI_LI | CHA | 271 | Late Intermediate | AD 900-1450 | Northern Peru |
|  | TOTAL |  | 1629 |  |  |  |

**Fig 1.**
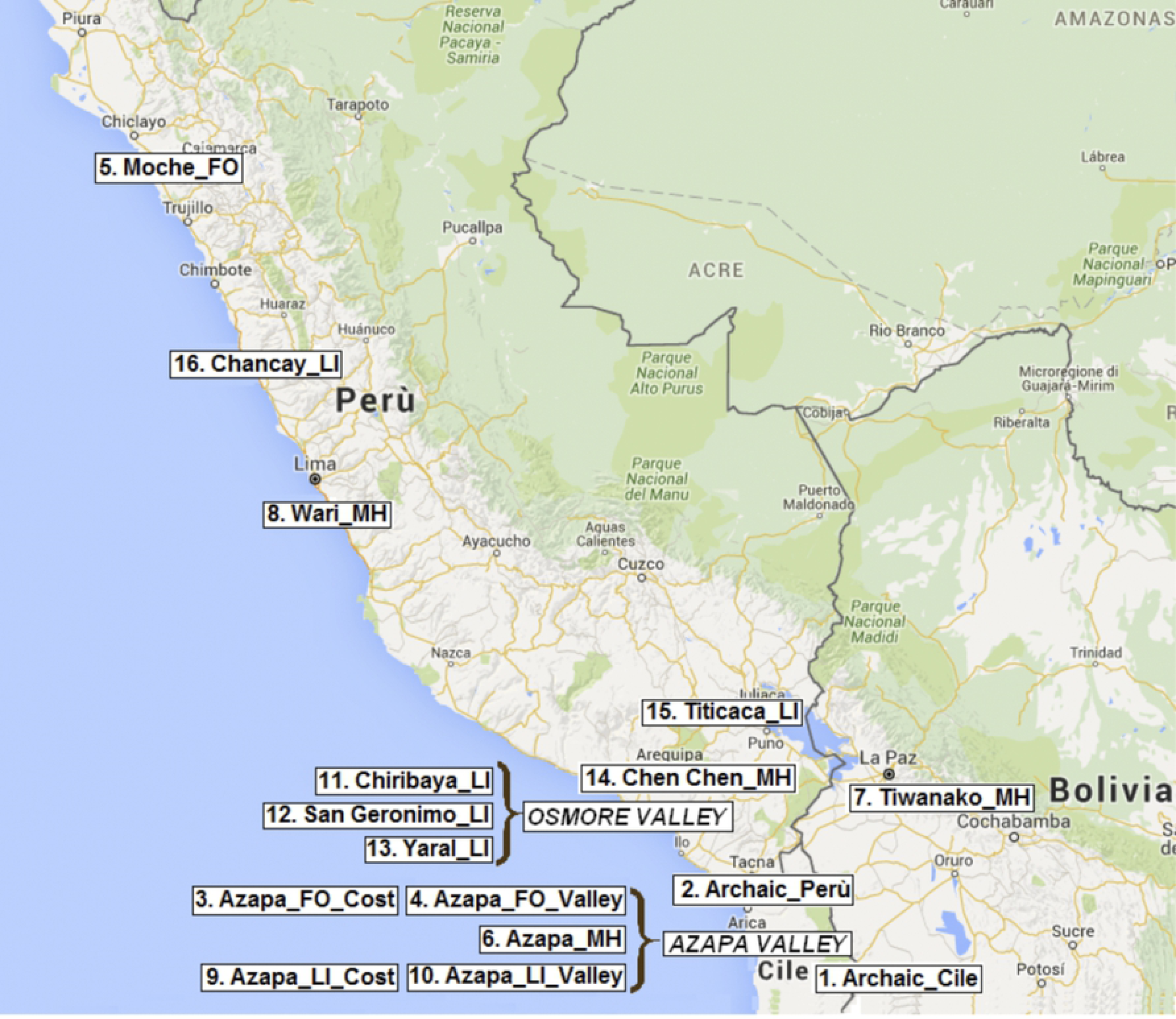
Location of the sixteen Pre-Inca sites from Peru, Bolivia and northern Chile included in the study.

Data were collected on all individuals regardless of age or sex – the absence of relevant sex dimorphism in dental morphological characteristics enables the pooling of all individuals in population studies [38-41] – using the Arizona State University Dental Anthropology System (ASUDAS) [41], establishing the degree of expression for each of the antimeres and pooling the results for each side. When the antimeres differed, the one with the highest degree of expression was chosen, as indicated in the individual count method of Turner and Scott [42] and recommended by Turner et al. [41] (as abovementioned, individual data are listed in S2 through S17 Tables). After the selection of breakpoints for each of the dental characteristics considered, the original categorical data were simplified into a presence/absence dichotomy and the frequencies calculated for each of the 16 sites considered in the study (Table S18).

Prior to the study the authors responsible for scoring data calibrated with one another in order to ensure an agreement of more than 90% for each trait [43], so that inter-observer error would not compromise results. For more details on interobserver error, see Nichol and Turner [44].

Forty-seven traits were initially scored; eight of them were discarded due to the extremely low variation between samples, reducing to 39 the number of traits used in this study (24 maxillary and 15 mandibular traits) for the 1665 individuals pooled into 16 groups on the basis of chronology and provenance (see S18 Table for the frequencies of traits according to their respective breaking points).

The biological similarities between samples were assessed through maximum likelihood (ML) and the reliability of the results appraised through bootstrap analysis. The results obtained were corroborated conducting principal component analysis (PCA), mean measure of divergence (MMD) with the Freeman-Tukey angular transformation for high (>0.95) or low (<0.05) trait frequencies [45-46], multi-dimensional scaling (MDS), and cluster analysis (CA) by means of Ward’s grouping method. Because multivariate tests can produce slightly different results that depend on the specific grouping method and algorithm used [39], the selection of multiple quantitative methods grants that affinities observed between samples in more than one elaboration are due to real similarities and not merely statistical artifacts.

## Results

The ML analysis, supported by bootstrap (Fig 2), divides the samples into two clusters positioned at the opposite ends of an unrooted tree and separated by a node that appears in 85% of the iterations. The first of these two clusters includes the samples from Chancay (LI), Moche (FO), Chen Chen (MH), Wari (LI) and Tiwanaku (MH) and separates, in 45% of the iterations, the last three of these samples from the rest of the cluster. The length of the branch connecting Tiwanaku and Wari does however urge caution in evaluating possible similitudes among the two.

**Fig 2.**
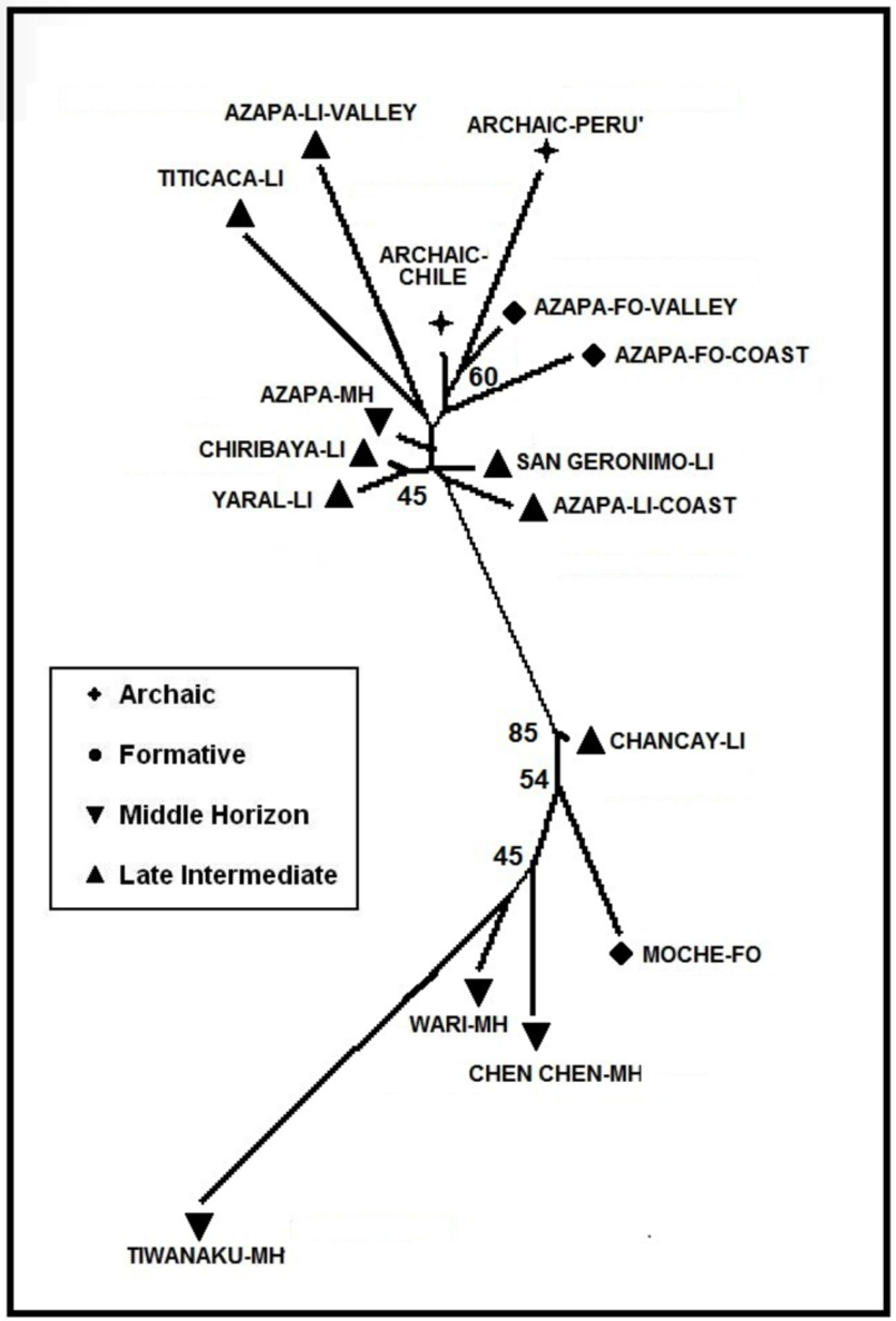
Maximum Likelihood unrooted tree with bootstrap values. Node values are expressed out of 100 iterations.

The second cluster includes all the Azapa Valley sites (coastal and inland), as well as the Chiribaya (LI), Yaral (LI) and San Geronimo (LI) samples from the Osmore Valley. The Titicaca sample from the region northwest of Lake Titicaca belongs to this second cluster and it shares a node with the Azapa-LI-Valley sample. In 60% of the iterations the more ancient samples included in this second cluster (Archaic Chile, Archaic-Peru, and the formative samples of the Azapa Valley) separate from all the more recent ones.

To corroborate the results obtained through ML we conducted a cluster analysis applying Ward’s grouping method (Fig 3). Similarly to what observed through ML, this too separates the samples into two main groups with Tiwanaku, Chen Chen, Wari, Moche and Chancay in one, and the remaining samples in the other.

**Fig 3.**
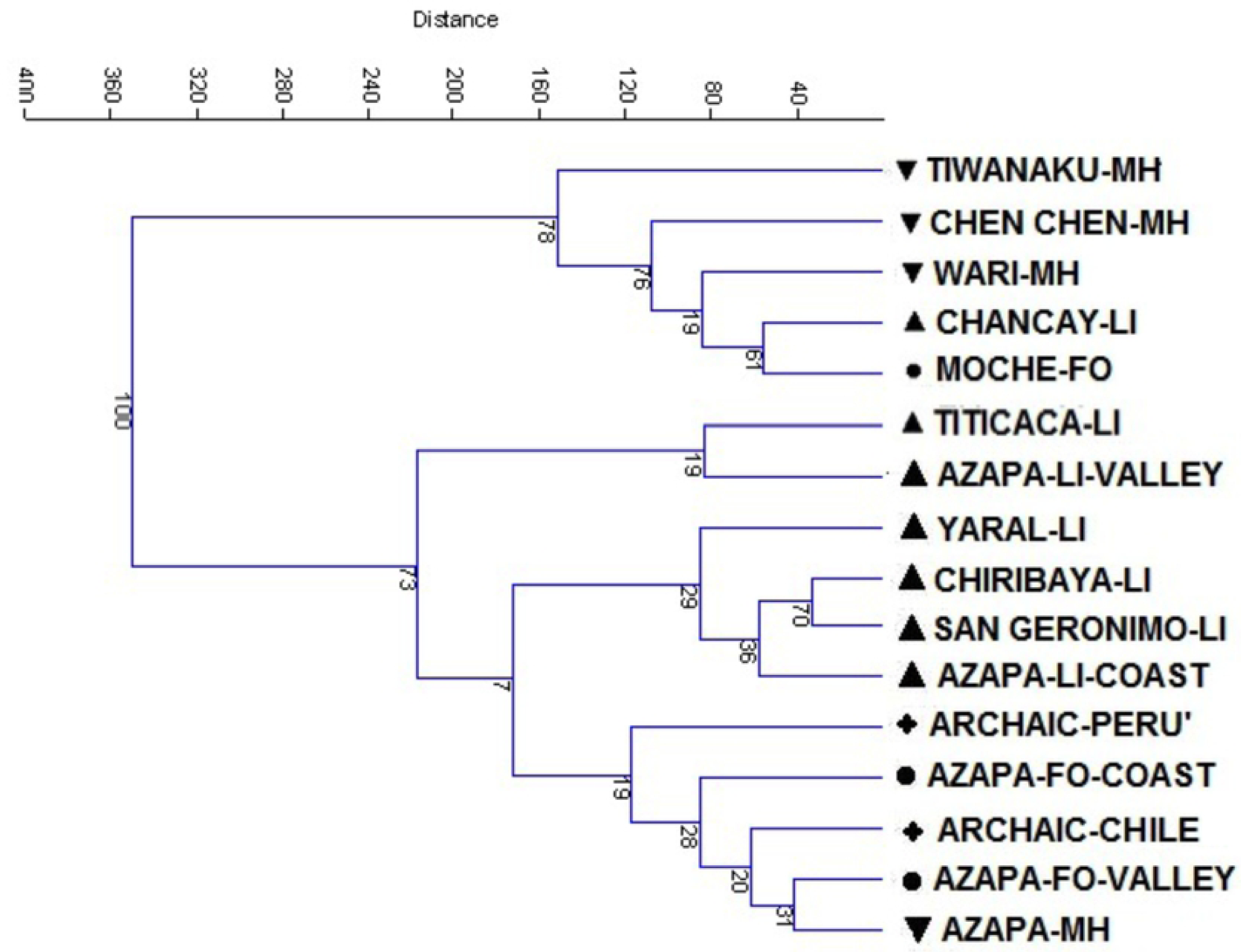
Cluster Analysis based on Ward’s grouping method. Node values are those obtained through 100 iterations.

Once again, Chen Chen shows similarities with Wari, Chancay and Moche. In 76% of the cases these form a cluster; Tiwanaku connects to this cluster through a long and independent branch in 78 out of 100 iterations. Of the remaining samples, the ones that appear to differ the most are Titicaca and Azapa-LI-Valley given that they separate from all the other samples of this second group in 73% of the iterations. The remaining samples form two sub-clusters; one comprises all the more ancient groups from Peru and Chile (Archaic and Formative), and the second the Late Intermediate coastal groups from both Chile and Peru. Caution must however be applied in drawing conclusions from these more detailed results given that bootstrap iterations are, in these cases, extremely low and all one may truly state regarding within cluster relations among samples is that there is some affinity between Chiribaya-LI and San Geronimo-LI given that these cluster in 70% of the iterations.

When applying principal component analysis (PCA) (Fig 4), Chen Chen, Wari, Tiwanaku, Moche and Chancay plot along the positive axis of the first component (which explains 28.8% of total variance); however, while Moche lays on the positive side of the second component (which explains about 14.2% of total variance), all the others set along its negative axis. Different traits concur in significantly influencing the distribution of the samples along the first axis, in which 10 traits present correlation coefficients above 0.7 (Table 2). All remaining samples plot along the first component’s negative axis and distribute in a manner that is, once again, in agreement with the trends observed in the previous analyses. Distribution reflects chronology to some extent, separating the more ancient (Archaic and Formative) samples from the more recent Middle Horizon and Late Intermediate ones. In this case the distribution is determined by traits that, in all but one case (lower M1 root number), have correlation coefficients that do not reach the established 0.7 thresholds.

**Table 2.**
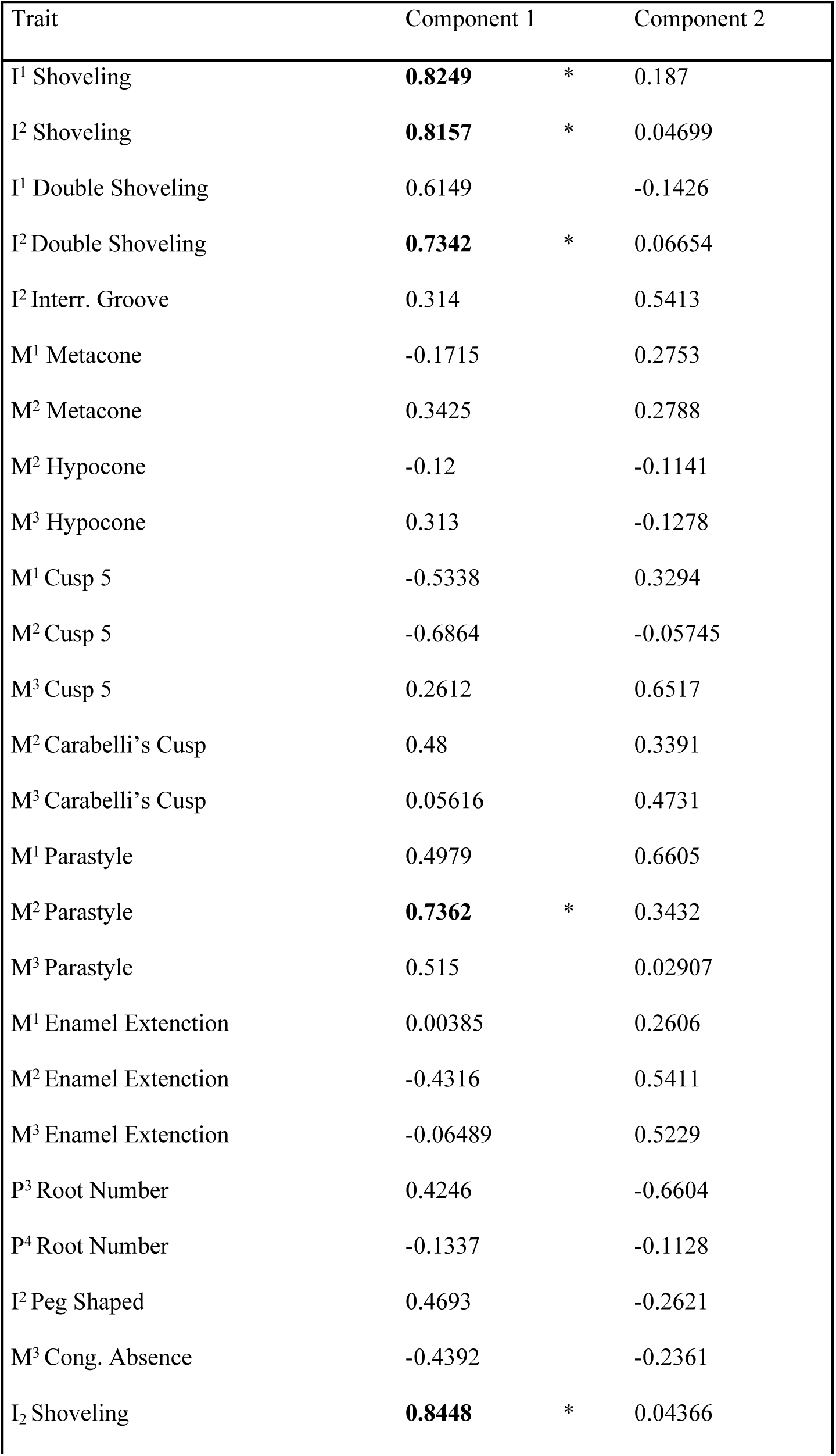
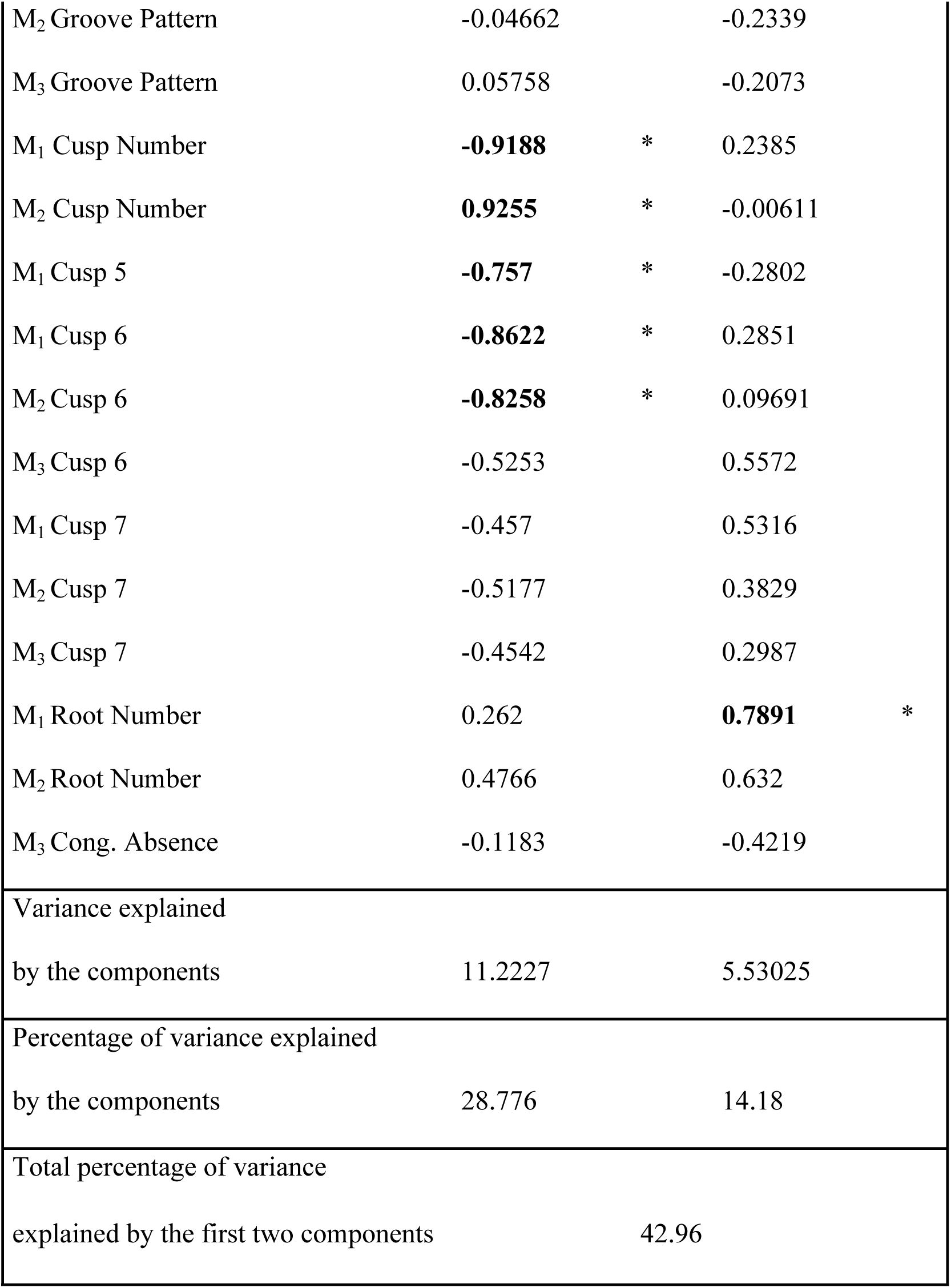
Principal Component Analysis factor loadings. Correlation coefficients greater than 0.7 are in bold and indicated with an asterisk.

**Fig 4.**
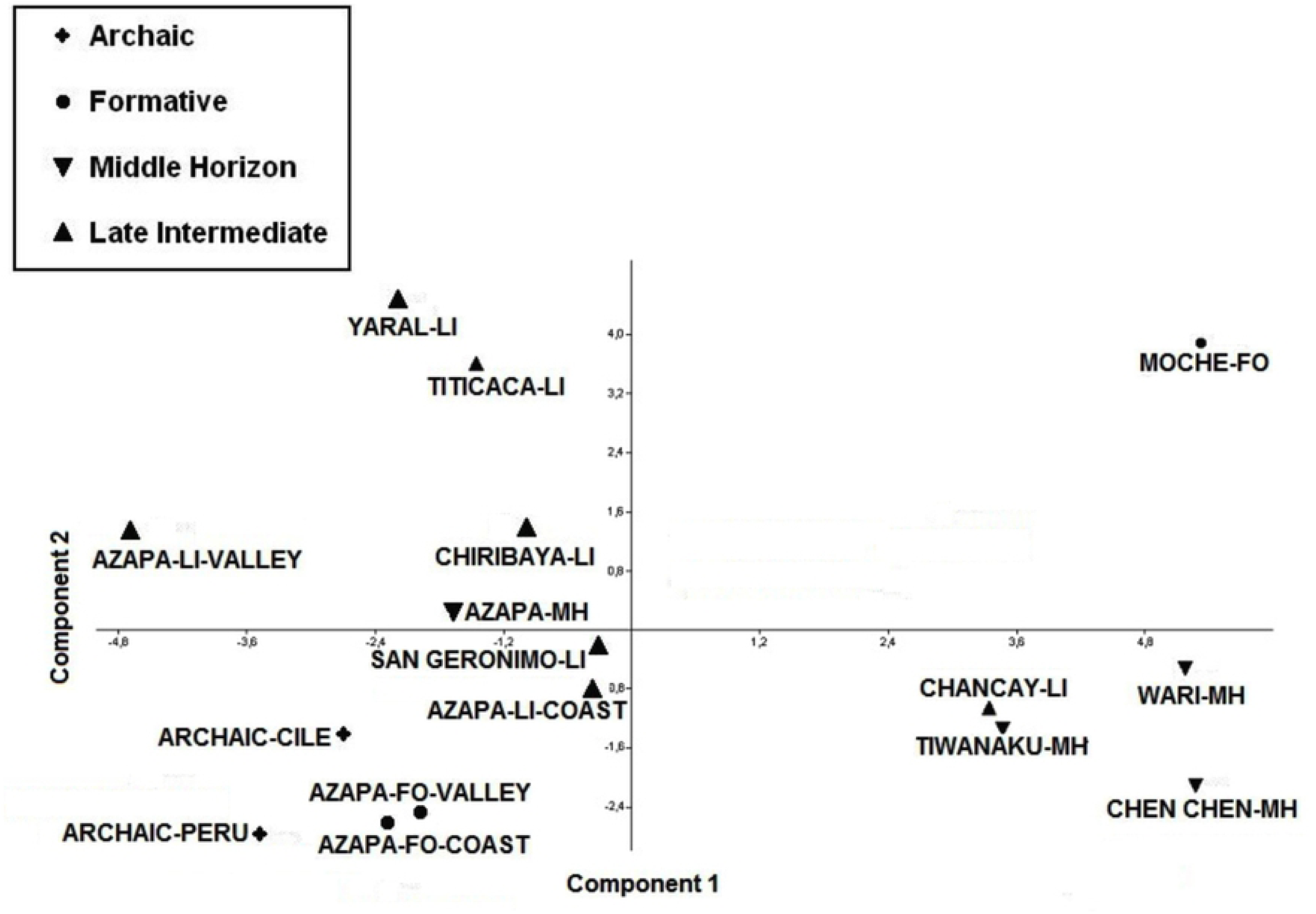
Bi-dimensional scatterplot from Principal Component Analysis.

Distance between groups was assessed through Mean Measure of Divergence (MMD) (Table 3). The highest MMD values (above 0.5, with the highest absolute value of 0.6240) obtained are those between Azapa-LI-Valley and the sites located in Peru (Chen Chen, Moche and Wari) and then those that separate Wari from the Titicaca Late Intermediate and from Yaral. In terms of noticeable distances, the Wari sample is the most distant one given that eight out of fifteen of the distances calculated are above 0.4. Most of these are with Azapa and Archaic groups, while the remaining ones are with San Geronimo, Titicaca and Yaral. Chen Chen is the next sample that separates the most from all other samples as six of the distances calculated are above 0.38. Similarly to what observed in the Wari sample, it too seems to differ most from the Azapa and Archaic groups and then from the Titicaca and Yaral ones.

Overall, the MMD matrix indicates that the Azapa and Archaic groups do not have remarkable intra-group distances, while they do show a fair amount of differentiation from the remaining samples. More varied and heterogeneous results can be observed in the other groups. Titicaca, Wari and Yaral show three distances above the 0.4 thresholds and San Geronimo two. Tiwanaku, Chiribaya and Chancay do not part noticeably from any group, despite the geographical (Chancay) or chronological barriers (Tiwanaku and Chiribaya). Interestingly, once again, Chen Chen, Wari and Tiwanaku do not show any noticeable distances among one another.

**Table 3.** Mean Measure of Divergence matrix. The values of divergence values are listed in the lower left portion, the corresponding standard deviations in the upper right one (bold values indicate statistically significant distances). Acronyms of sites are listed in Table 1.

## Discussion

The last decades have witnessed an intensification of bio-archaeological investigations on the ethnicity, population dynamics, cultural and genetic changes that characterized the Osmore Valley [5,15-19,21,23,26,47] and the Azapa Valley [2,13,20,22,24] in pre-Inca times. Although such studies have shed more light onto the relationships among Andean populations, and have contributed to confirm or reject contrasting hypotheses, several issues are still far from being solved.

From a geographical perspective, this study shows a separation between coastal and inland sites in Peru; at the same time it shows chronological continuity between the samples located in the coastal drainages of the Osmore Valley and the Azapa Valley. Owen [48] suggested that the collapse of the Tiwanaku state at the end of the Middle Horizon period led to a second diaspora of Tiwanaku colonists towards the coast, a continuity that found support in other studies [5,23,49-50]. On the contrary, Lozada Cerna [19,47] argued against this hypothesis suggesting that the Chiribaya groups didn’t have particular ties to the Middle Osmore Tiwanaku colonists. In this perspective, Sutter [2,22] found morphological differences between the coastal and the inland sites of the Azapa Valley; he also noted, on a larger scale, that they all clearly appeared to separate from the Late Intermediate Chiribaya, San Geronimo and El Yaral groups from the Coastal Osmore Valley. The differences observed with the Chiribaya groups [5] led him to interpret the Osmore Valley coastal drainage Chiribayas as a likely result of the Late Intermediate Tiwanaku diaspora [49].

The inclusion of additional groups, as in the case of the present study, enables the detection of other relations and large-scale trends. The similarities shown between the Archaic group from Peru and the Chilean groups from the Azapa Coast and Valley seem to clearly indicate there may have been chronological continuity with the formative period. At the same time, the fact that the Late Intermediate samples from Chiribaya, San Geronimo and El Yaral tend to cluster with the Late Intermediate coastal groups for the Azapa Valley and, even if to a lesser extent, to the Late Intermediate Azapa Valley group suggests this continuity might also have been geographical. Once again there appears to be a certain distinction between the coastal and inland sites of the Azapa Valley in both the formative and, even though somewhat less, the Late Intermediate. However the change in perspective compared to Sutter’s analysis [5] suggests this might denote less distance than originally proposed by his study. Chiribaya, San Geronimo and El Yaral do tend to form a sub-cluster that falls within a larger one that includes all the Azapa groups except for the Azapa-LI Coast one. While this is not at all that dissimilar from what was obtained in Sutter’s research from a regional perspective, the inclusion of further samples places these results in a new perspective in which, in light of the much greater distances obtained between the samples of Chiribaya, San Geronimo and El Yaral and those of the Osmore Valley, the distances obtained between these sites and the Azapa groups become less significant.

Biocultural markers such as cranial deformation patterns have suggested there may have been, in the Middle and Upper Osmore Valley, a close relationship between the Tiwanaku highland population and the one from their enclaves the valley (i.e., Chen Chen) [16-17]. Even though supported by archaeological material culture [3] and stable isotopes evidence – strontium and oxygen that indicate migration into the Osmore Valley of individuals from regions characterized by isotopic values compatible with those of the southern Titicaca basin [27,51] – these ties do not preclude there may have been others between Chen Chen and some of the nearby Wari outposts of the Middle Osmore Valley such as Cerro Baul. For example, evidence of this was found by us in a recent study conducted on a restricted portion of the samples used in the present analysis [18], in which we detected affinities between Chen Chen and both a Wari sample and, even though to a lesser extent, a Moche one. The relations detected among samples strongly suggested these ties had been anything but negligible but they likewise appear to indicate that, had isolation by cultural barriers occurred [52], as proposed by Goldstein [3], Chen Chen should have manifested a lower degree of similarity to those groups. Instead, the results seemed to suggest that, after the establishment of Cerro Baul and of the other outposts, Wari colonists must have necessarily come into direct contact with their Tiwanaku counterparts as also hypothesized by Moseley et al. [28]. The inclusion, in the present study, of the Tiwanaku sample from the southern Titicaca basin provides further evidence that the Wari and Tiwanaku colonists came into direct contact in the Middle Osmore Valley. In all the multivariate analyses conducted Chen Chen plots closer to the Middle Horizon Wari sample than it does to the Tiwanaku sample it ought to cluster with if indeed it were its ancestral population and one it continued to have ties to whilst maintaining isolation from the surrounding Wari populations. In general, the fact that the Tiwanaku collection clusters in different manners to the group formed by Chen Chen, Wari, Moche and Chancay, while it clearly diverges from Ciribaya, San Geronimo and El Yaral, as well as from all the sites (coastal and inland) in the Azapa Valley, suggests the existence of a general and widespread common morphological background for the inhabitants of the Middle Osmore Valley sites that is independent of their cultural affiliation.

The well-defined distance between coastal and inland samples in the Osmore Valley suggests the presence of a geographic or cultural barrier to the gene flow between these two regions. While the Chiribaya and San Geronimo sites are close to the coast, the site of El Yaral is located not far from the border between the coastal region and the Middle Osmore Valley. Yet, El Yaral clusters with the coastal samples and is morphologically distant from both Chen Chen and Tiwanaku. Overall, the relevant distances observed between Chiribaya, San Geronimo and El Yaral combined to those seen between Tiwanaku and Chen Chen appear to be more compatible with Lozada Cerna’s theory [19,47], according to which the Late Intermediate coastal sites in the Osmore drainage were independent from the Tiwanaku colonies, than it does with Owen’s [49] hypothesis of a second Tiwanaku diaspora. However, in this perspective we must maintain caution and view the results as the indicators they are and not as unequivocal ones, given the Tiwanaku sample dates to the Middle Horizon, while El Yaral, San Geronimo and Chiribaya to the Late Intermediate. The hypothesized second diaspora occurred at the onset of the Late Intermediate period, just after the fall of the Tiwanaku empire around AD 1000-1100 onward [8,53-55], as a likely consequence of climatic (drought) changes and the sociopolitical collapse in the *altiplano* [9,56-59].

Furthermore, the diaspora could have given rise to low-status groups that integrated, as minorities, in the coastal Osmore Valley populations [48] or, given the ethnical diversity attributed to the Tiwanaku population – described as characterized by independent “moieties, ethnic groups, or maximal ayllus” [3: p51] – we could simply have included Tiwanaku samples that were not representative of those who might have (potentially, if ever) migrated towards the coast.

There is little doubt that Tiwanaku and Wari represented two separate and independent empires, as the archaeological evidence shows [3]. Both of their outposts in the Osmore Valley produced ceramics, similar in style to those found in their respective heartlands, using locally available clay procured within their respective territories [60]. Even though the respective inhabitants of the enclaves of these two polities certainly imported cultural practices from their respective homelands, the present study reveals patterns of mobility and population dynamics within southern Peru and northern Chile that contrast with the generalized ideas of separation based on cultural identity.

The proximity shown in all analyses between Chen Chen and the Wari groups indicates that the interpretations based on cultural characteristics and separations might be insufficient to comprehend the complex peopling history of the region. Archaeological evidence [3] clearly assimilates Chen Chen to Tiwanaku and suggests minimal interaction and cultural assimilation between these two important, Tiwanaku and Wari, polities. The similarities detected in the present study, based on dental morphological characteristics, suggest Chen Chen had much stronger ties to Wari than indicated by cultural parameters alone thus denoting a certain degree of genetic homogeneity all around Peru as well as a paucity in morphological diversity. The same occurs with the Tiwanaku sample, which clusters with Wari, Moche and Chancay.

The patterns of population dynamics in Central-Southern Peru are also discussed by Knudson and Tung [33: p.307], who state that the presence of “non-local individuals at the site of Beringa derived from geologic zones located in the Pacific coastal areas, or what is now southern Peru, parts of Bolivia, or northern Chile, is consistent with the notion that the Middle Horizon was marked by expansive trade networks and population movement, likely influenced by local, historically durable trade practices and policies of the Wari empire”. This pattern of genetic homogeneity finds support in Kemp et al. [32], who suggested genetic continuity between Wari and post-Wari populations based on mtDNA analysis. Nonetheless, the authors do not rule out that the genetic continuity could be the result of an overall homogeneity in the region, which may mask (should it have occurred) possible female migrations into the Wari Empire after its collapse. Similarly, Lewis et al. [61] found that the mtDNA variation between the skeletal sample from Chen Chen and a large number of ancient and extant South American populations indicated strong haplogroup homogeneity in the region of southern Peru and northern Chile. They assessed that this homogeneity existed well before the Middle Horizon, the period that witnessed the rise of both the Wari empire and the Tiwanaku state. According to Lewis [62], variations in mtDNA, found among ancient and contemporary groups in the region, is more likely the result of genetic drift and not of genetic exchange (i.e., migrations) through time.

The Late Intermediate Titicaca sample also presents some peculiarities for its positioning in the plots does not appear to fit properly with any of the proposed models. Its positioning does however seem to find explanation when analyzed in light of genetic drift, isolation, and founder effect concepts. The Titicaca collection comes from the highlands long the northwestern side of the Lake Titicaca, at the opposite end of the Tiwanaku state and away from the sphere of influence of the Wari Empire. The distance shown between this group and all the other – geographically closer – groups in southern Peru, could consequently be the result of isolation by distance and genetic drift.

## Acknowledgements

We wish to thank all of the Museums and Institutions that made this study possible by granting us access to the collections and in particular (listed alphabetically): The American Museum of Natural History, New York, USA; the Anthropology Museum “S. Sergi”, Sapienza University, Rome, Italy; the Archaeological Museum, Universidad Nacional del Altiplano, Puno, Peru; the LCHES Duchworth Foundation, Cambridge University, UK; the Musée de l’Homme, Paris, France; and the Museo Contisuyo, Moquegua, Perù.

The exchange program for professors of the Ministry of Education, University and Research (MIUR) within the framework of the “IV Executive program of the Cultural Agreement between the Government of the Italian Republic and the Government of the Republic of Peru for the years 2002-2006”, which allowed to carry out several study missions in Peru. The final stage of this publication was in part supported by CONACyT grant CB-2017-2018-A1-S-10037. The present work has been furthermore supported by the PRIN Project (Ministero dell’Istruzione dell’Università e della Ricerca) “A multi-species genomic approach to assess pre- and post-Columbian population dynamics in South America” Grant N.: 20174BTC4R and the H2020 Programme ARIADNEplus project, contract no. H2020-INFRAIA-2018-1-823914. The authors opinion do not necessarily reflect those of the European Commission.

## Supporting information

**S1 Table. List of all the collections analyzed, their acronym, chronological period and location**

**S2 Table. Individual morphological data in the Archaic_Chile group.**

**S3 Table. Individual morphological data in the Archaic_Peru group.**

**S4 Table. Individual morphological data in the Azapa_Coast_Formative group.**

**S5 Table. Individual morphological data in the Azapa_Valley_Formative group.**

**S6 Table. Individual morphological data in the Moche_Formative group.**

**S7 Table. Individual morphological data in the Azapa_Middle_Horizon group.**

**S8 Table. Individual morphological data in the Tiwanaku_Middle_Horizon group.**

**S9 Table. Individual morphological data in the Wari_Middle_Horizon group.**

**S10 Table. Individual morphological data in the Azapa_Coast_Late_Intermediate group.**

**S11 Table. Individual morphological data in the Azapa_Valley_Late_Intermediate group.**

**S12 Table. Individual morphological data in the Chiribaya_Late_Intermediate group.**

**S13 Table. Individual morphological data in the San_Geronimo_Late_Intermediate group.**

**S14 Table. Individual morphological data in the Yaral_Late_Intermediate group.**

**S15 Table. Individual morphological data in the Chen_Chen_Late_Intermediate group.**

**S16 Table. Individual morphological data in the Titicaca_Late_Intermediate group.**

**S17 Table. Individual morphological data in the Chancay_Late_Intermediate group.**

**S18 Table. Frequency of the 39 traits used for statistical analyses in the 16 groups according to their own breaking points.**

## Data availability statement

Data was collected in a number of institutions and museums in France, Italy, Peru, the UK and the USA (listed alphabetically):

- American Museum of Natural History, New York, USA
- Anthropology Museum “S. Sergi”, Sapienza University, Rome, Italy
- Archaeological Museum, Universidad Nacional del Altiplano, Puno, Peru
- LCHES Duchworth Foundation, Cambridge University, UK
- Musée de l’Homme, Paris, France;
- Museo Contisuyo, Moquegua, Perù

